# Group B meningococcal outer membrane protein vaccine promote potent anti-viral effect

**DOI:** 10.1101/645457

**Authors:** Web Smith, John Smith

**Author notes:** Address: 631 Sumter Street, Columbia, SC 29208.

## Abstract

This report demonstrates a novel method to explore and evaluate the specific humoral/cellular immune response levels and immunoprotective effects of NMB0315 nucleic acid vaccine, recombinant protein vaccine and nucleic acid vaccine + recombinant protein vaccine in combination with mice, and to further explore the effective immunization method for NMB0315 vaccine. This route provides experimental basis. Nucleic acid vaccines [pcDNA3 1(+) / NMB0315] and recombinant protein vaccines (pET 30a / NMB0315) were prepared in large quantities, and immunologically or separately immunized female BALB/c mice were determined by nucleic acid priming protein boosting method. The specific humoral/cell immune response level, the in vitro bactericidal titer of immune serum, and the immunoprotective effect of the vaccine on mice infected with group B meningococcus were observed. Serum-specific IgG, IgG1, IgG2a and genital lavage fluids induced by NMB0315 nucleic acid vaccine group (pNMB0315 CpG), protein vaccine group (rNMB0315 FA) and combined immunization group (pNMB0315 CpG+rNMB0315 FA). The specific sIgA level reached the peak in the eighth week, and the A450 values were in vitro, and the in vitro bactericidal antibody titers of the nucleic acid vaccine group, the protein vaccine group and the combined immunization group were 1, 64, 1128, respectively. The immune protection rate of experimental mice were 70%, 95% and 80%, respectively. At 2, 4, 6, and 8 weeks, the ratio of IgG2a / IgG1 in the nucleic acid vaccine group, the recombinant protein vaccine group, and the combined immunization vaccine group was less than 1.

## Introduction

Neisseria meningitides (Nm) is a Gram-negative diplococcus that is obligately parasitic to humans, also known as meningococcus. In bacterial meningitis, epidemic cerebrospinal meningitis (flowing brain) caused by meningococcal disease is the only disease that causes an outbreak. The bacteria can invade the brain and spinal cord of humans, causing suppurative infections, which can cause permanent damage to the brain and nervous system [1 4]. Nm’s Capsular polysaccharide (CPS), Outer membrane protein (OMP) and lipooligosaccharides are important virulence factors [5]. At present, the C group, A+C group and the mixed vaccines of group A, C, W and Y developed by capsular polysaccharide prion protein carrier effectively control meningococcal diseases caused by group A, C, W and Y. [6]. The CPS immunogenicity of the group B meningococcus serogroup B (MenB) is weak, and the structure is similar to the structure of N-acetylacetylneuraminic acid in human neural tissue, which may cause cross-reaction and trigger autoimmune diseases, limiting B. The successful development of the cerebral capsular polysaccharide vaccine [7,8]. Epidemic outbreaks are a global public health problem. According to statistics, about 50% of meningococcal diseases worldwide are caused by MenB [9]. Therefore, the development of an effective B-group epidemic vaccine has far-reaching social significance and broad clinical application prospects. Group B Neisseria meningitides serogroup B0315 (NMB0315) is an outer membrane protein of MenB consisting of 430 amino acids with a coding gene of about 1 293 bp and a molecular weight of 46 kD [10]. In 2011, Wang et al. [10] studied the crystal structure of NMB0315 and found that the antigenicity of the carboxy-terminal domain of the outer membrane protein is highly conserved. Blast analysis revealed that it has a different serogroup between meningococcal bacteria and Neisseria gonorrhoeae. The homology reached 98%. Description NMB0315 can be used as a potential candidate vaccine antigen for group B meningococcus and can be used as a universal vaccine candidate antigen for other serogroups. Studies have shown that nucleic acid vaccines induce strong cellular immune responses while protein vaccines induce stronger humoral immune responses. In order to improve the vaccine immune effect, the researchers used a combination of different types of vaccines to play a different immune response induced by different vaccines; through the complementary advantages, synergy and other effects to induce the body to produce a high level of specific immune response, thereby Improve the immune effect of the vaccine [11,12]. Based on the NMB0315 nucleic acid vaccine and recombinant protein vaccine constructed by the research group [13,14], the NMB0315 nucleic acid vaccine priming recombinant protein vaccine was used to enhance the female BALB / c via muscle and intraperitoneal route respectively. In rats, it is preliminary to explore whether the effect of DNA prime protein boost combined immunization is better than that of nucleic acid vaccine alone or protein vaccine alone, which provides an experimental basis for further exploring the effective immunization methods and pathways of NMB0315 vaccine.

## Materials and Method

Group B Neisseria meningitidis standard MC58 was purchased from ATCC, USA; E coli BL21 recombinant expression strain containing pET 30a / NMB0315 and E coil JM109 recombinant clone of pcDNA3 1 (+) / NMB0315 (referred to as pNMB0315) All of them were constructed and preserved by our group; 3 to 4 weeks of female BALB / c mice were purchased from Hunan Slack Jingda Experimental Animal Co., Ltd.; anti-NMB0315 serum was obtained by immunizing New Zealand rabbits; HRP-labeled goat anti-rabbit IgG, restricted Dicer (BamH I, Xho I), T4 ligase and protein Marker were purchased from Fermentas, the plasmid was purchased from Omega, and the adjuvant CpG was synthesized by Shanghai Biotech. The sequence was 5 TCTCACCGTTCCTGACGTT 3 Model 1826, full chain sulfation modification; Freund adjuvant was purchased from Sigma, and ELISA kit was purchased from eBioscience.

Preparation of protein vaccine and nucleic acid vaccine The recombinant cloned E coilJM109 containing pcDNA3 1 (+) and pNMB0315 was amplified and cultured, and the plasmid was digested with BamH and Xho, and the endotoxin plasmid extraction reagent was removed. The eukaryotic plasmid was extracted in large quantities, and the concentration of the extracted plasmid was detected by a nucleic acid protein analyzer. The E coli BL21 expressing bacteria containing the prokaryotic plasmid pET 30a / NMB0315 were amplified and cultured, and the recombinant protein NMB0315 was expressed by Ni Zhen Li Zhenyu and other group B meningococcal outer membrane protein 0315. 10 phase · 1503 · NTA affinity chromatography purification, BCA method for protein concentration determination, SDS PAGE analysis of protein purity, −80 benefits for storage.

Eighty BA 3 / 4 female BALB / c female mice were randomly divided into three groups: A, B, and C (PBS group, nucleic acid vaccine pNMB0315 + CpG group and rNMB0315+FA group, FA was Freund’s adjuvant). And one experimental group D (co-immunization group: pNMB0315 CpG+rNMB0315 FA), 20 mice per group. Group B intramuscular injection of nucleic acid vaccine 50 ng each time, group C intraperitoneal injection of protein vaccine 30 ng each time, each group was immunized 4 times at 0, 2, 4, 6 weeks; group D 0, 2 weeks for nucleic acid vaccine muscle Note that 4 and 6 weeks were intraperitoneal injections of protein vaccines. At the same time, blood was collected from the tail vein of the mouse at 0, 2, 4, and 6 weeks, and vaginal secretions were collected with a standard sterile cotton swab. The 80% refrigerator was frozen; after 2 weeks of the last immunization, the mouse eyeball was taken in the biosafety cabinet. Blood, the mice were sacrificed and the spleen was aseptically obtained, and 200 mesh nylon gauze was ground and filtered to prepare a single cell suspension.

The humoral immune response level was detected by using the purified recombinant protein NMB0315 as an antigen-coated ELISA plate, mouse genital lavage or serum as the primary antibody, and indirect ELISA to detect total serum IgG and subclass, genital tract lavage in mice. Liquid sIgA level.

Detection of cellular immune response levels The proliferation of spleen lymphocytes in immunized mice was detected by CCK 8 colorimetric assay. After 2 weeks of the last immunization, the spleens of the mice were aseptically isolated to prepare a lymphocyte suspension, and the cell concentration was adjusted to 6 0 6 10 ml −1, and added to a 96-well plate, 100 μl / L well; 20 stimuli, 3 replicate wells were set, and medium without recombinant protein was used as a blank control. Place 5% CO2, 37 y for 48 h, add 10 mM l / L CCK 8 solution, 5% CO2, 37 s for 4 h, and determine the absorbance at 450 nm (A) by microplate reader. Stimulation index (SI) = (A stimuli - A blank) / (A unstimulated well - A blank). The level of IFN in the supernatant was determined by ELISA.

Two weeks after the last immunization, the mice were challenged with a lethal dose (40 000 CFU/ml) of Neisseria meningitidis MC58, and the incidence and mortality of the mice were observed within 72 hours, and the survival rate was calculated [15]. The MC58 broth that grows to the log phase is mixed with the rabbit rabbit 1 1, and then added to the test serum at a ratio of 1 2 (56 inactivated, diluted by dilution), fully mixed and then cultured for 37 h. Add chocolate agar medium, mix and post 37 liters overnight, and set a positive control (diagnostic serum) and a negative control. The negative control includes bacterial liquid plus complement, the bacterial liquid heats the inactivated complement, and the serum plus bacteria heats up. Live complement three holes. When the results are judged, the reciprocal of the highest dilution of serum with a sterilization rate of 50% or more is based on the complement negative control, which is the serum antibody bactericidal titer [15].

The experimental data was analyzed with statistical software SPSS16 0. Antibody levels, spleen cell stimulation index and cytokine levels were expressed as x s, and the mean between groups was analyzed by analysis of variance, P < 0 05, which was statistically significant.

## Results and Discussion

Identification of Nucleic Acid Vaccines and Protein Vaccines The eukaryotic recombinant plasmid pNMB0315 retained by our group was digested with BamH jade and/or Xho jade to obtain a 1 293 bp target band (Fig. 1). The purified recombinant plasmid pET 30a / NMB0315 was analyzed by Western blot and a clear specific target band was observed at about 46 kD (Fig. 2).

**Figure 1.**
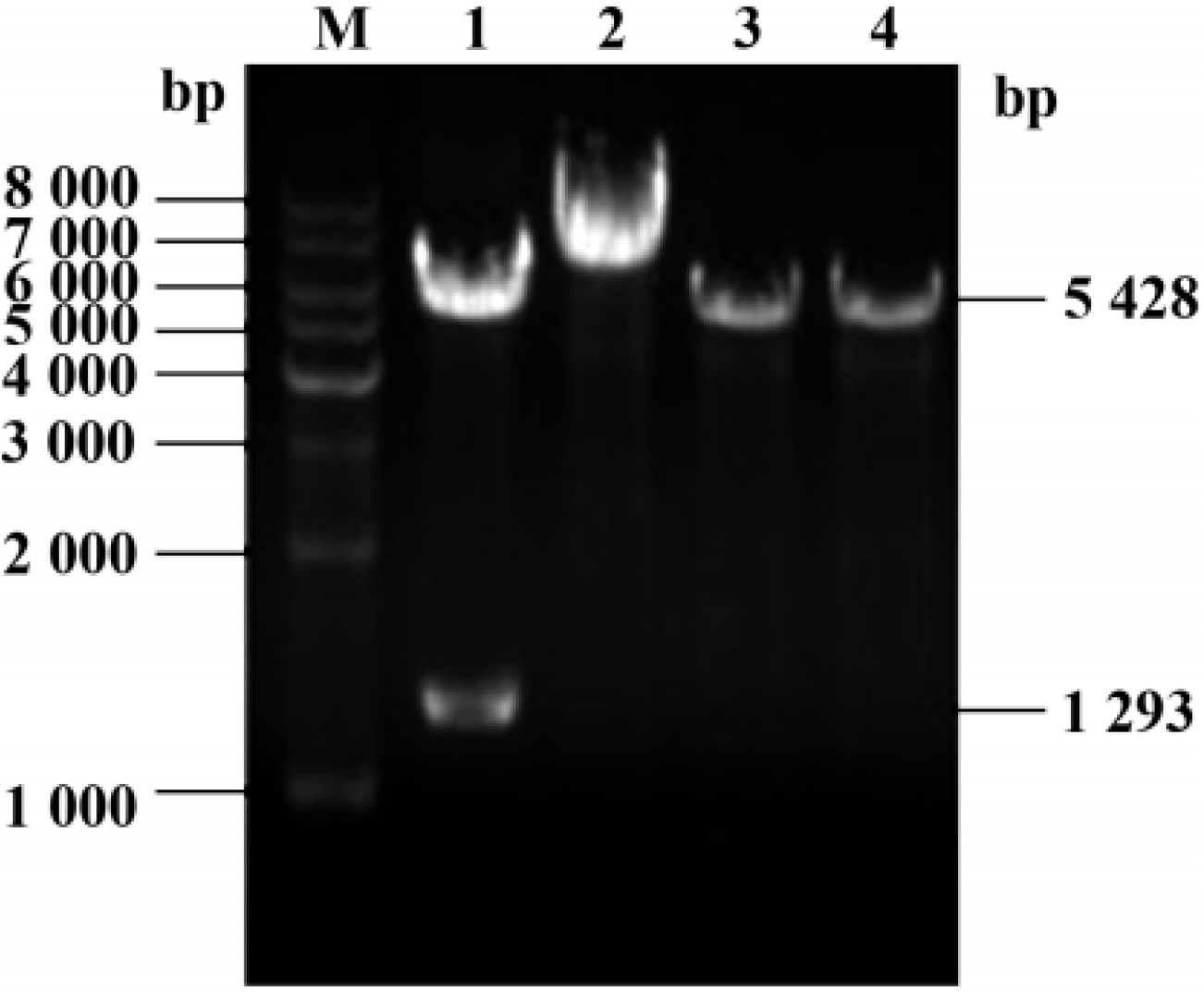
Gel electrophoresis

**Figure 2.**
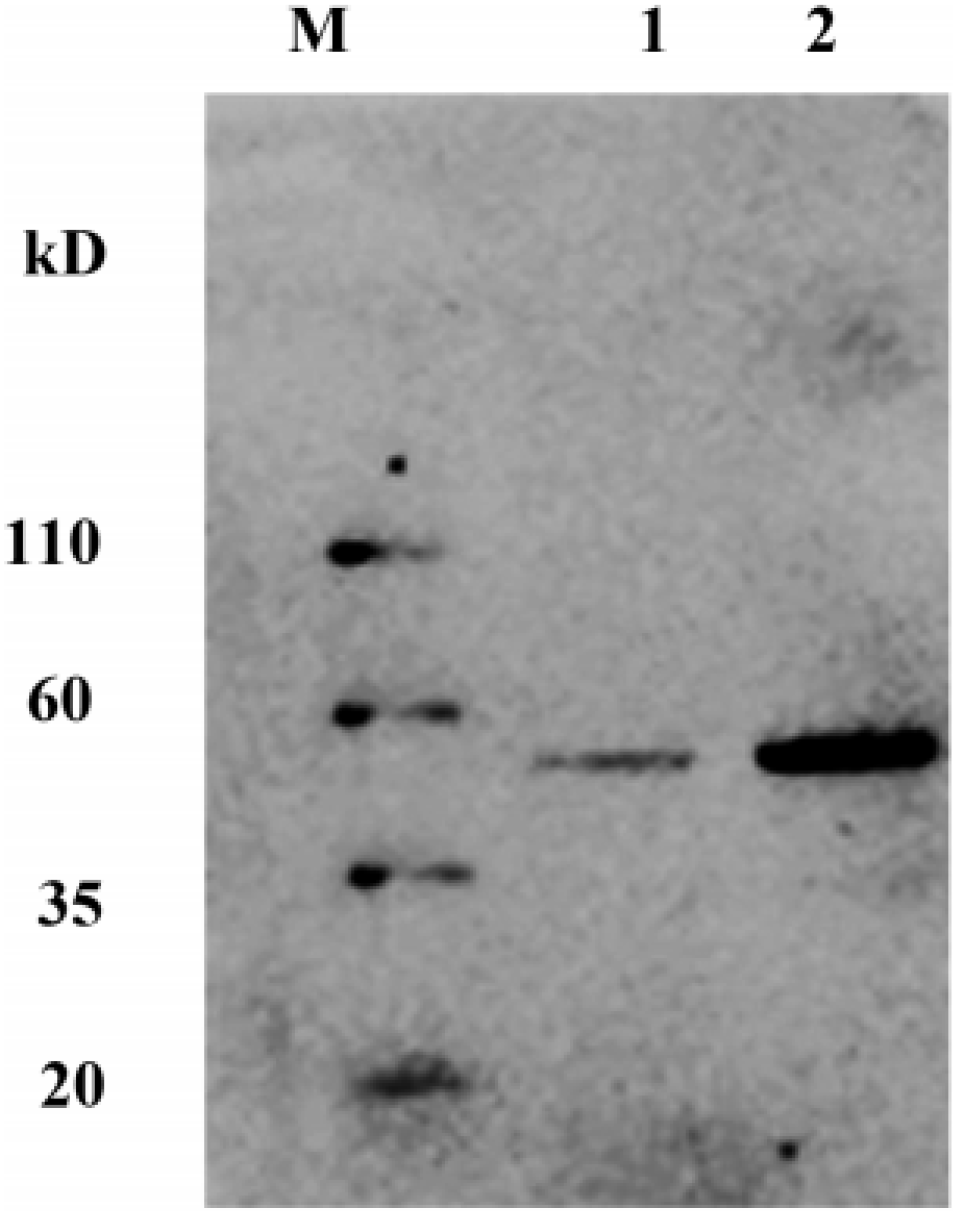
Western blot

**Figure 3.**
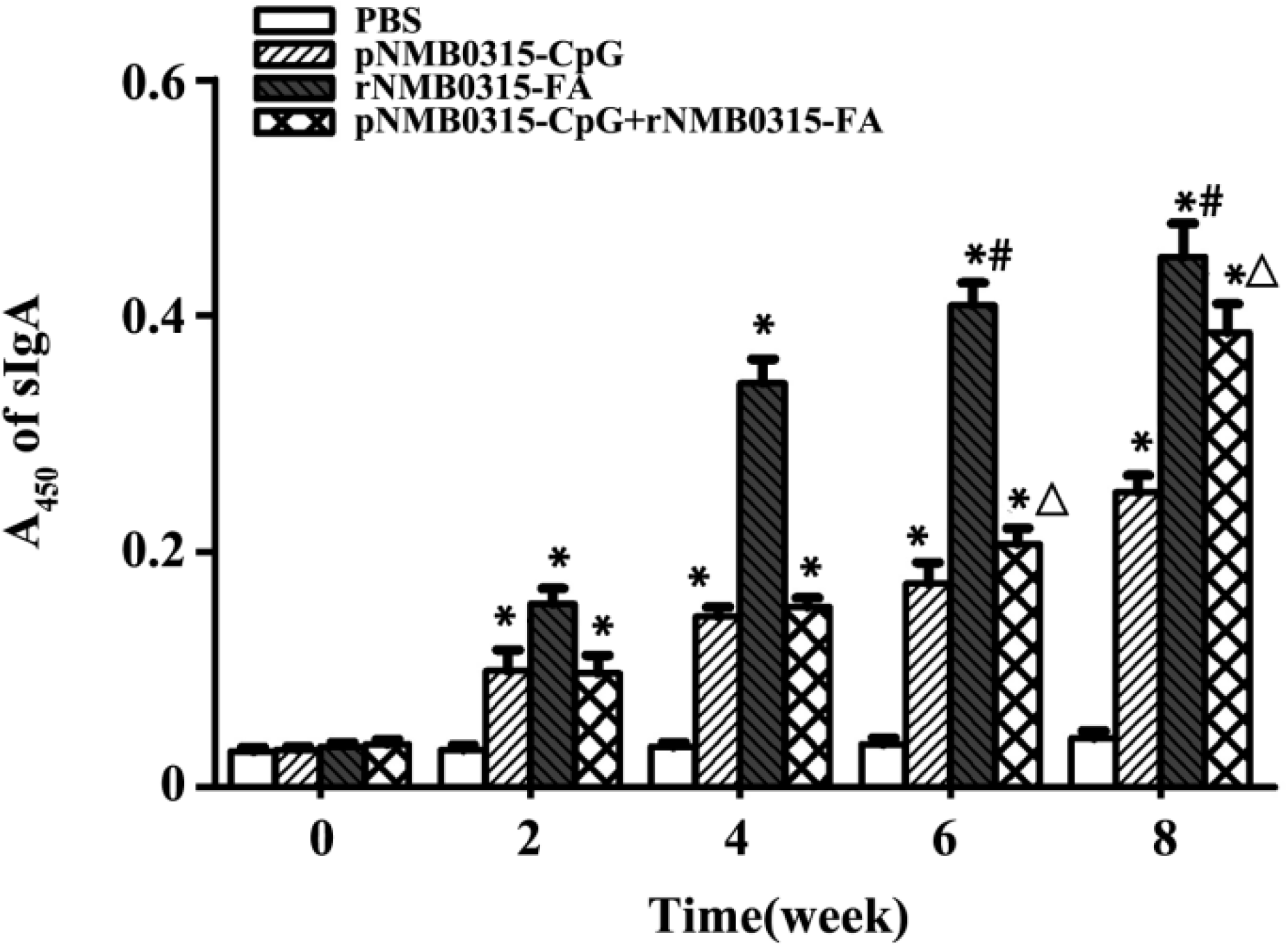
ELISA analysis

Mouse genital mucosa-specific sIgA levels were collected from mouse genital mucosal secretions at 0, 2, 4, 6, and 8 weeks. Recombinant protein rNMB0315 was used as antigen to coat ELISA plates, and specific sIgA levels were detected by indirect ELISA. The results showed that with the increase of immunization time, the levels of sIgA in genital lavage fluid of group B (pNMB0315 CpG), group C (rNMB0315 FA), group D (pNMB0315 CpG+rNMB0315 FA) increased. At the 8th week, the peaks were reached. The A450s in groups B, C, and D were (0 250 to 0 015), (0 450 to 0 028), and (0 386 to 0 024), which was significantly higher. Group A.

Immunologically specific IgG levels were obtained from the immunized mice. The specific IgG levels in the serum of the immunized mice were detected by indirect ELISA using rNMB0315 as the coating antigen. The results showed that serum IgG levels in group B (pNMB0315 CpG), group C (rNMB0315 FA), group D (pNMB0315 CpG + rNMB0315 FA) increased significantly with increasing immunization time, peaking at week 8 The A450s in groups B, C, and D were (0 505 according to 0 016), (0 823 by 0 019), and (0 694 by 0 013), which were significantly higher than group A (PBS control group, P < 0 01), and the level of group C is the highest, which is significantly higher than that of group B and D (P < 0 05). The results are shown in Figure 4.

**Figure 4.**
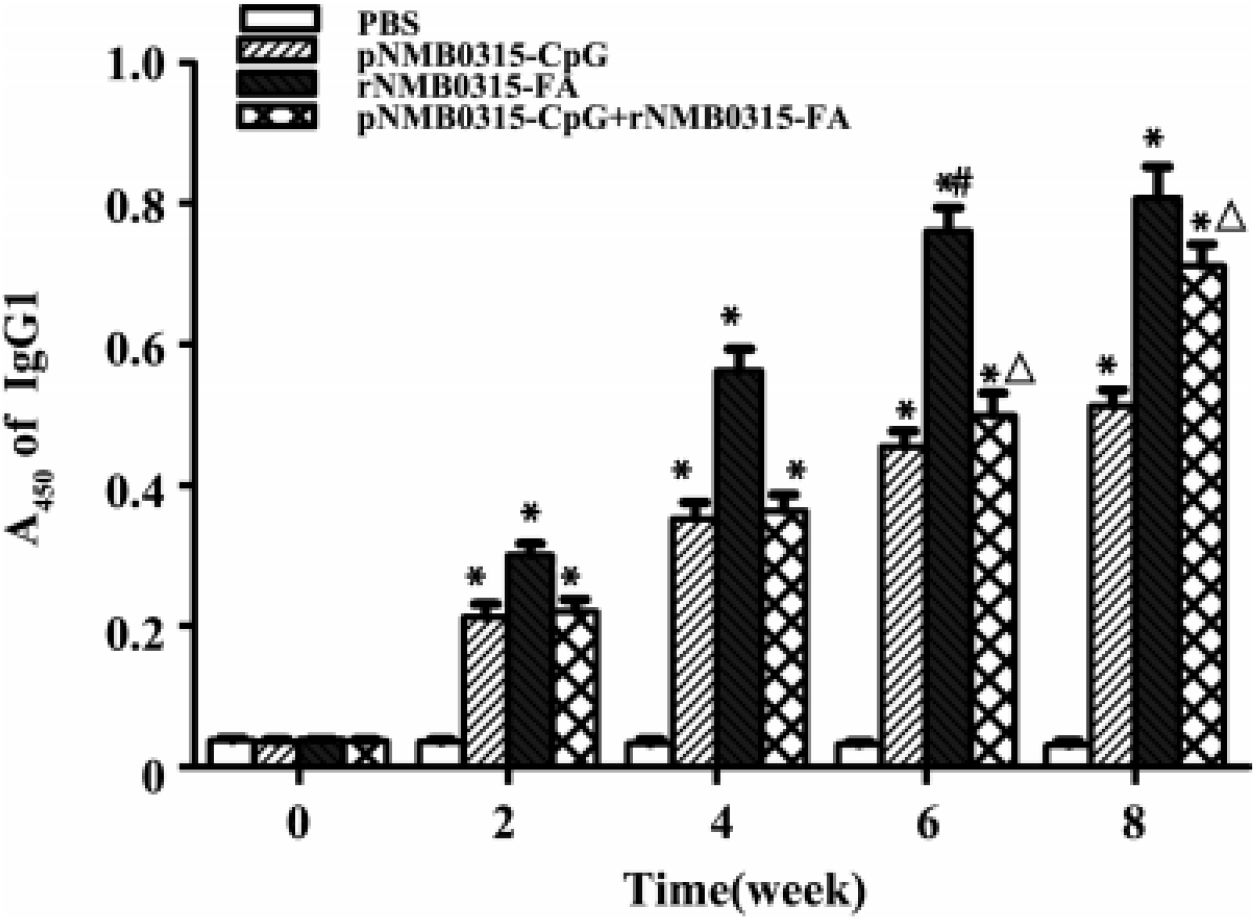
Antibody titer analysis

The detection method is the same as the above IgG. The results showed that the serum levels of IgG1 in group B (pNMB0315 CpG), group C (rNMB0315 FA), group D (pNMB0315 CpG+rNMB0315 FA) increased with the number of immunizations, peaked at week 8, A450. The values were (0 513 according to 0 022), (0 807 to 0 045) and (0 711 to 0 032), and there was a significant difference compared with the PBS group (P < 0 01). At week 6 and week 8, the level of specific IgG1 in serum of group C was significantly higher than that of group B and group D.

The ratio of IgG2a / IgG1 in the pNMB0315 CpG group was 0 741 (0 158 / 0 213), 0 731 (0) when the specific IgG2a / IgG1 ratio in the immunized mice was immunized for 2, 4, 6 and 8 weeks. 258 / 0 353), 0 695 (0 316 / 0 455) and 0 667 (0 342 / 0 513); rNMB0315 FA group IgG2a / IgG1 ratio is 0 771 (0 232 / 0 301), 0 716 (0 403 / 0 563), 0 714 (0 543 / 0 761) and 0 739 (0 596 / 0 807); pNMB 0315 The ratio of IgG2a / IgG1 in the CpG+ rNMB0315 FA group was 0 665 (0 147 / 0 221), 0 712 (0 259 / 0 364), 0 695 (0 347 / 0 499) and 0 640 (0 455 / 0 711), the ratio of IgG2a / IgG1 in each group was less than 1.

Immune mouse-specific cellular immunity levels Mice spleen lymphocytes were stimulated with recombinant protein NMB0315, and the spleen lymphocyte stimulation index (SI) of each group was measured by CCK8 colorimetry. The results showed that the SI values of spleen lymphocytes in the pNMB0315 CpG group, rNMB0315 FA group and pNMB0315 CpG+rNMB0315 FA group were 1 846 depending on 0 143, 1 476±0 060, 2 185°0. 086) was significantly higher than the PBS group (P < 0 01); the SI value of the pNMB0315 CpG + rNMB0315 FA group was significantly higher than that of the pNMB0315 CpG and rNMB0315 FA groups (P nucleic acid vaccine group>protein vaccine group).

The spleen lymphocyte supernatant of the immunized mice was subjected to IFN concentration for two weeks. The IFN assay was used to detect the IFN concentration in the spleen lymphocyte supernatant obtained by the recombinant protein stimulation. The results showed that: pNMB0315 In the CpG group, rNMB0315 FA and pNMB0315 CpG+rNMB0315 FA group, the spleen lymphocyte culture supernatant of the immunized mice had a IFN concentration of 170 85 to 18 85 pg/ml and 120 77 to 14 respectively. 70 pg / ml and 226 60 were significantly higher than 22 76 pg / ml in the PBS control group (P < 0 01); and the pNMB0315 CpG+ rNMB0315 FA group was higher than the pNMB0315 CpG and rNMB0315 FA groups (P nucleic acid vaccine group> protein vaccine group, see Figure 8.

Immune protection effect of vaccine and in vitro bactericidal activity of immune serum Two weeks after the last immunization, the lethal dose of Neisseria meningitidis MC58 was used to attack the mice, and the morbidity and mortality of the mice were observed every day, and the survival rate was calculated. The results showed that the survival rates of group B (pNMB0315 CpG), group C (rNMB0315 FA), group D (pN MB0315 CpG+ rNMB0315 FA) were 70%, 95% and 80%, respectively, which were significantly higher than PBS group. (PFA) > co-immunization group (pNMB0315 CpG + rNMB0315 FA)>nucleic acid vaccine group (pNMB0315 CpG). When B, C, and D immune sera were diluted to 1 64, 1 128, and 1 128, respectively, more than 50% of the experimental bacteria MC58 died, ie, groups B, C, and D, under the complement-mediated effect. The in vitro bactericidal antibody titers of the mouse immune serum were 1 64, 1 128 and 1 128, respectively.

Nucleic acid vaccines and recombinant protein vaccines are widely used in vaccine research against pathogens to induce specific immune responses. In order to improve the vaccine immune effect, the researchers tried to use different types of vaccine combined immunization strategies to play different immune responses induced by different vaccines [11]. The combination of nucleic acid priming protein-enhanced immunoassay has made progress in the study of vaccines such as schistosomiasis, Japanese encephalitis, brucellosis, AIDS, tuberculosis^1–3^, malaria, etc., and its immune response is superior to a single nucleic acid vaccine or Protein vaccine [16 19]. Whether the N. meningitidis NMB0315 vaccine adopts the nucleic acid priming protein-enhanced combination immunization method is superior to the protein vaccine or nucleic acid vaccine alone^4–12^, has not been reported yet, and deserves further research and exploration.

In this study, NMB0315 nucleic acid vaccine and recombinant protein vaccine were used to immunize female BALB/c mice by muscle and intraperitoneal route respectively, and the immune effect of DNA prime protein boost combined with immunization was preliminarily investigated. The results showed that the nucleic acid vaccine group (pNMB0315 CpG), the recombinant protein vaccine group (rNMB0315 FA) and the combined immunization group (pNMB0315 CpG + rNMB0315 FA) induced specific IgG, IgG1, IgG2a and sIgA antibody levels in mice. It showed an upward trend with time and peaked in the eighth week^13–20^. The antibody titers of the nucleic acid vaccine group were 1 40 000, 1 50 000, 1 40 000 and 1 25,000 respectively. The antibody titers of the recombinant protein vaccine group were respectively 1 19 000, 1 90 000, 1 80 000 and 1 35,000; the antibody titers of the combined immunization group were 1 150 000, 1 70 000, 1 50 000 and 1 30 000, respectively. They were significantly higher than the PBS control group (P combined immunization group > nucleic acid vaccine group. Meanwhile, the immune protection effect of NMB0315 vaccine on experimental mice (Yi 14 d) was 95% in the protein vaccine group, 80% in the combined immunization group, and nucleic acid. The vaccine group was 70%; the antibody titers of the serum bactericidal assay (SBA) were 1 128 for the protein vaccine group, 1 128 for the combined immunization group, and 1 64 for the nucleic acid vaccine group. Neisseria meningitidis (Nm) For extracellular parasites, anti-Nm infection mainly relies on humoral immunity. [20] These three groups further demonstrate that the recombinant vaccination group with good humoral immunity has in vitro bactericidal effect and immune protection effect on experimental mice. It is also the best. In this study, nucleic acid priming prion protein was used to enhance the combined immunization method, and the induced humoral immunity and cellular immunity level were higher than the single nucleic acid vaccine group, but the combined immunity group induced humoral immunity level was lower than that of single recombinant protein vaccine. The reason may be that the NMB0315 nucleic acid vaccine is not efficient in transfection in mouse muscle cells and the expression level of effective immunogen is limited. The IgG subclass in serum can roughly reflect the change of the body’s immune response type. The Th2 predisposing humoral immune response is predominant, and IgG2a is predominantly reflecting the Th1 propensity for cellular immune response. [21] At the 8th week of vaccine immunization, the serum-specific IgG2a / in the nucleic acid vaccine group, the recombinant protein vaccine group, and the co-immunized group mice. The ratio of IgG1 is 0 667 (0 342 / 0 513), 0 739 (0 596 / 0 807) 640 0 Yan (Yan 455 0/0 Yan 711), a ratio less than 1. Studies have found that IFN secreted by Th1 promotes IgG2a production, thereby promoting cellular immune responses and T-killer cell activation. The combined vaccination group induced IFN levels significantly higher than the nucleic acid vaccine group and the protein vaccine group.

Based on the NMB0315 nucleic acid vaccine and recombinant protein vaccine constructed by the research team, NMB0315 nucleic acid vaccine and recombinant protein vaccine were used to jointly immunize female BALB/c mice through muscle and intraperitoneal route respectively. Whether the effect of DNA prime protein boost combined immunization is superior to a single nucleic acid vaccine or a separate protein vaccine. The results showed that the humoral immunity induced by the recombinant protein vaccine and the immunoprotection effect on the experimental mice were the best, followed by the combined immunization with the priming of the nucleic acid. The above results can provide experimental basis for further exploration of effective immunization methods and pathways of NMB0315 vaccine, and lay the experimental foundation for the development of high-efficiency B group influenza vaccine.

## Conclusion

The effects of NMB0315 vaccine-induced humoral immunity (including mucosal immunity) are as follows: recombinant protein vaccine group, combined immunization vaccine group, and nucleic acid vaccine group; the effect of inducing cellular immunity from high to low is: combined immunization vaccine group, nucleic acid vaccine Group, recombinant protein vaccine group; NMB0315 vaccine immune test effect on experimental mice from high to low: recombinant protein vaccine group, combined immunization vaccine group, nucleic acid vaccine group.

